# Genetic structure in the wood mouse and the bank vole: contrasting patterns in a human-modified and highly fragmented landscape

**DOI:** 10.1101/464057

**Authors:** Roberto Biello, Andrea Brunelli, Giulia Sozio, Katja Havenstein, Alessio Mortelliti, Valerio Ketmaier, Giorgio Bertorelle

**Author notes:** Corresponding authors: Giorgio Bertorelle,; Roberto Biello.

## Abstract

Habitat fragmentation related to human activities modifies the distribution and the demographic trajectory of a species, often leading to genetic erosion and increased extinction risks. Understanding the impact of fragmentation on different species that co-exist in the same area becomes extremely important. Here we estimated the impact produced by different natural and anthropic landscape features on gene flow patterns in two sympatric species sampled in the same locations. Our main goal was to identify shared and private factors in the comparison among species. 199 bank voles and 194 wood mice were collected in 15 woodlands in a fragmented landscape, and genotyped at 8 and 7 microsatellites, respectively. Genetic variation and structure were analysed with standard approaches. Effective migration surfaces, isolation by resistance analysis, and regression with randomization were used to study isolation by distance and to estimate the relative importance of land cover elements on gene flow. Genetic structure was similarly affected by isolation by distance in these species, but the isolation-by-resistance analysis suggests that i) the wood mouse has constrained patterns of dispersal across woodland patches and facilitated connectivity in cultivated areas; ii) the bank vole connectivity is hindered by urban areas, while permeability is facilitated by the presence of woodlands, and cultivated terrains. Habitat loss and fragmentation can therefore influence genetic structure of small sympatric mammal species in different ways, and predicting the genetic consequences of these events using only one species may be misleading.

## Introduction

Habitat loss and fragmentation have negative impacts on populations, and are considered as one of the main causes of biodiversity loss and therefore a major issue in conservation biology^1–3^. In particular, anthropogenic habitat fragmentation has modified the distribution and population sizes in many different organisms^4,5^, with local and/or global reduction of genetic diversity and connectivity^6,7^. Monitoring the genetic consequences of human activities that increase habitat fragmentation is therefore important to develop appropriate conservation and management strategies^8^.

The major consequence of habitat loss and fragmentation is to create discontinuities (i.e. patchiness) in the distribution of critical resources (e.g. food, cover, water) or in environmental conditions (e.g. microclimate)^9^. Such discontinuities reduce connectivity among populations^10^, threatening their long-term viability due to genetic (e.g., reduced evolutionary potential and inbreeding depression) and demographic factors (e.g. demographic stochasticity)^11^. Habitat fragmentation may also have different short term consequences in different species, for example by reducing the suitable habitats or increasing the predation success, but these effects poorly predict long-term responses^12^. Gene flow among subpopulations is necessary to alleviate the adverse genetic consequences of population fragmentation, reducing genetic drift and maintaining local genetic variation^13^. From a conservation perspective, inferring the functional connectivity of populations across landscapes becomes crucial^9,14^. Identifying the areas where gene flow is either facilitated or prevented, and the landscape factors responsible for that, is a high priority^15,16^.

One interesting opportunity to investigate the causes and the genetic consequences of fragmentation is represented by sympatric species with partially overlapped ecological niches^17–19^. Different species, in fact, may respond very differently to the same landscape matrix^20–23^. They may also react differently to the fragmentation of their previously continuous habitat, and these differences may be reflected in the geographic distribution of their genetic variation. In this work, we investigate the effects of habitat fragmentation present in agricultural landscape in Central Italy on the genetic structure of two sympatric rodent species, the wood mouse (*Apodemus sylvaticus*) and the bank vole (*Myodes glareolus*).

The wood mouse is a generalist species known to inhabit a wide range of habitats including forests, hedgerows and agricultural fields^24–26^. In contrast, the bank vole is a “forest specialist”, i.e. it is more strictly associated with forest habitats, from mature stands to recently coppiced woodlands^27,28^. In general, specialist species tend to be more affected than generalist species by habitat fragmentation, both because highly dispersed resources are more difficult to reach by the former^29–31^, but also because of competitive exclusion of the specialists by the generalists^32^. Accordingly, the specialist bank vole seems to prefer sites with high connectivity^32,33^, and the generalist wood mouse can also be found in highly fragmented habitats, being able for example to move across cultivated fields^32,34^. We currently do not known whether these differences directly correspond to a stronger genetic structure in the bank vole compared to the wood mouse, and if (and how) different natural or anthropogenic habitat features have different relative impacts on gene flow. Our study aims at investigating these questions following three steps: (1) initially, neutral genetic markers will be used to estimate the genetic diversity and the population structure separately in each species; (2) patterns of gene flow and the geographic location of genetic barriers will be then analysed in the two species and compared; (3) finally, species-specific landscape features with the largest influence on the genetic variation pattern will be identified.

## Materials and Methods

### Study area and sample method

The study was conducted in a fragmented landscape (<20% of residual woodland cover) located in central Italy (coordinates: 42°30’50”, 12°4’40”; elevation: 350 m; Fig. 1). Woodland patches, consisting of mixed deciduous forest dominated by downy and turkey oaks *(Quercus pubescens* and *Quercus cerris*, respectively), were embedded in an agricultural matrix (mainly wheat fields) crossed by a network of hedgerows providing structural connectivity to habitat patches. The S2 highway and the railway bisect the study area, potentially acting as barriers to wildlife movements^35^. Finally, urban areas are present and represent approximately 5% of the total area. Twelve trapping sessions were conducted over a 2-year period, with trapping taking place every other month from April 2011 to February 2013. During each session, grids were trapped for three consecutive nights. Total sample size was 199 for the bank voles and 194 for the wood mice, and samples sizes in each of 15 different woodland patches is reported in Table 1. All the procedures of trapping and manipulation of animals took place in compliance with the European Council Directive 92/43EEC (Italian law D.Lgs 157/92 and LR 3/1994) and with the European Council Directive 86/609/EEC (Italian law D.Lgs 116/92). The capture and handling of species listed in the EU Habitat Directive was covered by permit number PNM 0024822 granted to A. M. by the Ministry of Environment, Rome, Italy.

**Figure 1.**
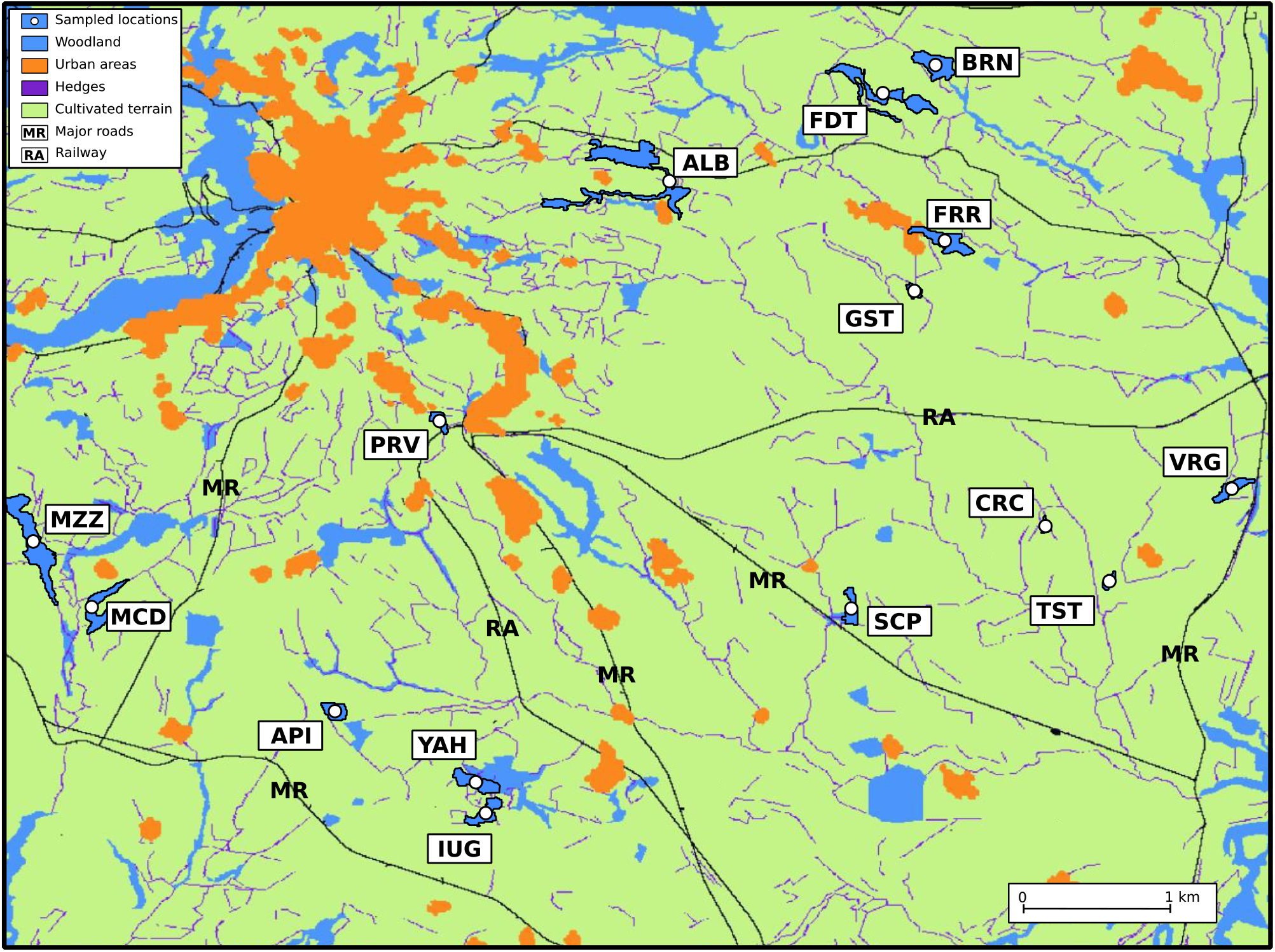
The study area. It is located in the Province of Viterbo, Central Italy. Landscape is reclassified according to the features utilized in the IBR analysis. RA represent the only railway intersecting the study area. Population codes as in Table 1.

**Table 1.** Genetic diversity indices in the wood mouse and the bank vole populations: sample size (N), number of alleles (Na), allelic richness (Ar), observed heterozygosity (Ho), and expected heterozygosity (He).

|  | *Wood mouse* |  |  |  |  | *Bank vole* |  |  |  |  |
| --- | --- | --- | --- | --- | --- | --- | --- | --- | --- | --- |
|  | N | Na | Ar | Ho | He | N | Na | Ar | Ho | He |
| ALB | 10 | 54 | 5.5 | 0.75 | 0.71 | 13 | 57 | 6.2 | 0.57 | 0.78 |
| BRN | 7 | 48 | 5.5 | 0.72 | 0.75 | 14 | 64 | 6.7 | 0.67 | 0.81 |
| FDT | 14 | 79 | 5.8 | 0.74 | 0.81 | 14 | 65 | 6.8 | 0.71 | 0.80 |
| FRR | 14 | 72 | 5.4 | 0.72 | 0.74 | 13 | 62 | 6.5 | 0.68 | 0.79 |
| GST | 14 | 62 | 4.8 | 0.66 | 0.75 | 14 | 57 | 5.9 | 0.72 | 0.76 |
| API | 14 | 66 | 5.0 | 0.73 | 0.71 | 14 | 47 | 5.2 | 0.72 | 0.74 |
| IUG | 14 | 65 | 5.0 | 0.70 | 0.73 | 14 | 52 | 5.8 | 0.67 | 0.77 |
| MCD | 14 | 65 | 5.2 | 0.64 | 0.69 | 14 | 44 | 4.8 | 0.65 | 0.68 |
| MZZ | 14 | 69 | 5.0 | 0.71 | 0.70 | 13 | 55 | 6.2 | 0.69 | 0.77 |
| PRV | 9 | 34 | 4.3 | 0.91 | 0.69 | 11 | 40 | 5.0 | 0.66 | 0.65 |
| YAH | 14 | 71 | 5.4 | 0.72 | 0.73 | 12 | 54 | 6.3 | 0.68 | 0.71 |
| CRC | 14 | 57 | 4.6 | 0.55 | 0.66 | 14 | 45 | 4.8 | 0.57 | 0.68 |
| SCP | 14 | 66 | 5.1 | 0.69 | 0.73 | 13 | 57 | 6.3 | 0.56 | 0.77 |
| TST | 14 | 65 | 5.1 | 0.66 | 0.68 | 13 | 50 | 5.6 | 0.59 | 0.69 |
| VRG | 14 | 65 | 5.1 | 0.66 | 0.73 | 13 | 49 | 5.7 | 0.67 | 0.76 |
| Mean | 12.9 | 62.5 | 5.1 | 0.71 | 0.72 | 13.3 | 53.2 | 5.9 | 0.65 | 0.74 |

### Genotyping

Genomic DNA was extracted from the mouse ear lobe samples using the NucleoSpin^®^ Tissue (Macherey-Nagel, Düren, Germany) according to the manufacturer’s protocol or using the Chelex-based DNA extraction method^36^. Eight microsatellite loci were used for the bank vole: Cg13B8, Cg6A1, Cg3F12, Cg13H9, Cg2E2, Cg3E10, Cg2A4 and Cg3A8^37^. Seven microsatellite loci, described for members of the genus *Apodemus*, were used for the wood mouse: As-7, As11, As-12, As-20, As-34, GTTD9A and MsAf-8^38–40^. A two-step PCR with the following conditions was carried out: initial denaturation at 95°C for 15 minutes, followed by 30 cycles at 95°C for 30 seconds, 56°C for 45 seconds and 72°C for 45 seconds, followed by eight cycles at 95°C for 30 seconds, 53°C for 45 seconds and 72°C for 45 seconds, and a final elongation at 72°C for 30 minutes. The forward primers were 5 labelled with one of the following fluorescent labels: FAM, VIC, NED and PET. Fragments were analysed on an ABI3130 capillary analyser (Applied Biosystems, Life Technologies Corporation). Fragment data were analysed using Peak Scanner Software (Applied Biosystems, Life Technologies Corporation).

### Genetic diversity

Descriptive statistics of nuclear genetic diversity were estimated separately for each population (woodland patch) in each species. The mean number of alleles, and the observed and expected heterozygosities, were estimated using Genalex 6.4^41^, and the same program was used to test for deviation from Hardy–Weinberg equilibrium. Allelic richness (AR) was calculated using the rarefaction procedure in the Fstat 2.9.3.2 software^42^. Arlequin 3.5.2.2^43^ was used to test for linkage disequilibrium between each pair of loci for each sampling population following a likelihood-ratio statistic, whose null distribution was obtained by a permutation procedure. We applied sequential Bonferroni corrections to account for multiple comparisons^44^. Micro-Checker 2.2.3^45^ was used to check for null alleles and scoring errors. FREENA^46^ was used to compare uncorrected and corrected F_ST_ values to test for the impact of null alleles, when present. Genetic differentiation measured as F_ST_ values^47^ was estimated for each pair of sampling population with Arlequin. Statistical significance of the F_ST_ values was tested using 10,000 permutations, and P values were multiplied by the total number of comparison following the conservative Bonferroni approach for multiple testing.

### Genetic structure

Two Bayesian clustering methods were used to identify the number of genetic groups without (STRUCTURE v2.3.4)^48^ and with (TESS v2.3.1)^49^ spatially explicit data. For the STRUCTURE analysis, a burn-in length of 50,000 iterations and a run length of 100,000 iterations were used in an admixture model with correlated allele frequencies among populations testing each K value between 1 and 15. Each K value was run 10 times. The optimal K value was determined using the ∆K method^50^ by means of STRUCTURE Harvester^51^. To visualize STRUCTURE results, STRUCTURE Harvester was used as well. CLUMPP^52^ was then applied to average the multiple runs given by STRUCTURE and to verify correct label switching. To display the results, the output from CLUMPP was visualized with DISTRUCT^53^. The CAR admixture model was used in TESS, with simple Euclidean geographic distances. We run 50,000 MCMC iterations with 20,000 burn-in for 12 times for each K value (2–15). We used deviance information criterion (DIC) values and stabilization of the Q-matrix of posterior probabilities to define the ideal number of clusters (i.e. K max) for the data (Ortego et al. 2015).

### Visualizing deviation from Isolation by Distance

Genetic diversity between populations often exhibit patterns consistent with Isolation by Distance (IBD)^55^, where populations far apart in the geographic space receive less gene flow than neighbouring ones. Given the ubiquity of this phenomenon^56,57^ it is interesting to see locations where this does not hold true, as they might represent barriers or zones of high contact. Global deviation from Isolation by Distance can be identified, for example, studying the decrease of similarity or autocorrelation with geographic distance. However, specific deviations in some areas, but not in others, cannot be easily investigated and visualized by standard methods. One recent answer to this problem comes from the use of Estimated Effective Migration Surfaces (EEMs)^58^. EEMS employs individual based migration rates in order to visualize zones with higher or lower migration with respect to the overall rate. These areas represent locations in which the pattern of gene flow predicted by IBD is facilitated or hindered. The region under study was first divided in a grid of demes and the individuals were assigned to the deme closest to their sampling location. The matrix of effective migration rates was then computed by EEMS based on the stepping-stone model^59^ and on resistance distances^60^. We used the EEMS script for microsatellites analysis runems_sats available from Github at https://github.com/dipetkov/eems to construct EEMS surfaces for the bank vole and the wood mouse. Considering that the number of demes simulated during the grid construction phase can influence the scale of the deviation from the overall migration rate, we averaged three runs with 50, 100, 200, 300 and 400 demes to produce the final EEMS surface. Each single run consisted in 200,000 burn in steps followed by 1,000,000 MCMC iterations sampled every 10,000 steps. We plotted the averaged EEMS and checked for MCMC convergence using the rEEMSplots package in R v 3.2.2.

### Isolation by resistance

Understanding the effect of environmental components on the genetic makeup of natural populations is the goal of landscape genetics, which integrates population genetics, landscape ecology and spatial statistics^61–63^. One of the techniques more commonly used in landscape genetics to identify discontinuities in gene flow and determine the relative resistance to movement imposed by different landscape elements is IBR, Isolation by Resistance^60^. IBR offers a conceptual model in which landscape resistance is the analogue of electrical resistance, and the movements of individuals and flow of genes are analogues of electrical current^64^. It greatly extends the ability to model multiple complementary paths of connectivity, while being sufficiently computationally efficient to allow its use over large landscapes at relatively fine resolution^65,66^. In order to analyse the effect of specific landscape components on gene flow, we tested for the presence of IBR. We first constructed a raster grid encompassing all our study area reclassifying the land cover based on features that were *a priori* most likely to affect gene flow in both the bank vole and the wood mouse: woodland, urban areas, cultivated terrain and hedges (Fig. 1). We also included in our raster grid the major roads intersecting our study area from OpenStreetMap (OpenStreetMap contributors, 2015) and the railways tracks from the DIVA-GIS database at http://www.diva-gis.org/gdatahttp://www.diva-gis.org/gdata.

In order to determine the relative importance of land cover elements in hindering or facilitating gene flow, we modified this grid under two different set of scenarios. The first set (resistance set) was aimed at determining the resistance caused by a specific land cover feature with respect to the others. We assigned a varying maximum resistance (RE_max_) to a target component, keeping the other landscape features to a uniform minimum resistance (RE_min_ = 1). The second set of grids (permeability set) was built to establish the possible role of a specific landscape feature in facilitating the connection between different populations. We assigned a minimum resistance value to a target landscape component and a varying RE_max_ to all remaining feature. For both set of grids we employed eight maximum resistance values (RE_max_ = 5, 10, 50, 100, 500, 1000, 5000 and 10000) obtaining a total of 96 different surfaces. We computed pairwise resistance distances between populations for both the bank vole and the wood mouse using the different sets of grids. Distances were obtained considering the eight-neighbour cell connection scheme in CIRCUITSCAPE 4.0^67^ with the sampled woodland patches as focal regions. We also computed an Isolation by Distance scenario considering a homogeneous resistance surface (all RE = 1)^54,68^. We then compared the resistance and the F_ST_ matrices using multiple matrix regression with randomization (MMRR)^69^. For each landscape variable, the most supported model was identified as the one corresponding to the highest supported *R^2^* value. In case of plateau, we preferred the model corresponding to the onset of the plateau^68^. Statistical significance of the coefficients was determined using 9999 permutations with the *MMRR* function^69^. Finally, for each species, we created a cumulative resistance surface assigning to every land cover variable the ratio of resistance with respect to RE_max_ obtained considering both set of models. We compared the output of CIRCUITSCAPE for these two cumulative grids with the F_ST_ matrix using MMRR and, to disentangle the effect of landscape features on genetic diversity from simple IBD, we computed a partial mantel test using the function *mantel.partial* from the package *vegan* version 2.4-2^70^. All statistical analyses were conducted in R v.3.2.2 (R Core Team 2016).

## Results

### Genetic diversity

All loci were polymorphic in both species. The average expected heterozygosities were very similar in the two different sets of markers typed in the two species (0.74 in the bank vole and 0.72 in the wood mouse), and the number of alleles varied between 2 and 16 in the wood mouse and between 3 and 11 in the bank vole markers, respectively. All the genetic variation statistics are reported in Table 1. No systematic deviation from linkage equilibrium was observed between loci for any population in both species, and none of the tests was significant after Bonferroni correction. Some loci showed evidence of the presence of null alleles, but only in some populations. We analysed the effect of these alleles by comparing matrices of pairwise F_ST_ values computed from the complete data set with values corrected for null alleles as estimated by FreeNA. Multilocus global F_ST_ values had identical values when calculated with and without correcting for null alleles in both species (wood mouse: F_ST_ = 0.03; bank vole: F_ST_ = 0.08), with identical or very similar confidence intervals in the two analyses (0.01–0.05 in wood mouse, with and without correction, 0.07–0.09 and 0.06–0.08 in bank vole, with and without correction, respectively). Multilocus pairwise F_ST_ values with and without correction were also highly correlated (wood mouse: *r* = 0.99; p = 0.001; bank vole: *r* = 0.99; p = 0.001; Mantel test). We decided therefore to use the complete data set for all downstream analyses. Pairwise F_ST_ values in the wood mouse were significant after sequential Bonferroni correction only in 7 out of 105 comparisons, all involving the PRV population (with F_ST_ values never larger than 0.08). On the contrary, the bank vole shows a much larger geographic structure. Approximately half of the F_ST_ values were significant, with the highest divergence values observed in comparisons including PRV, and, as reported above, the average F_ST_ was much higher than that estimated in the wood mouse.

### Genetic structure

The most likely partition implied three genetic groups (K=3) in both species. Here we present individual assignment plots for K equal to 2, 3 and 4 (Fig. 2A-B) to better visualize different aspects of the genetic structure, and we also report the geographic distribution of the most supported number of K in both species (Fig 2C). In the wood mouse (Fig. 2A), the isolation of PRV already suggested by the pairwise F_ST_ matrix was supported at different values of K. With the most supported K=3, or with K=4, a large fraction of individuals and populations (with the exception of PRV) showed a mixed ancestry. In the bank vole (Fig. 2B), populations appeared more internally homogeneous, with three distinct genetic groups prevailing in the northern areas (ALB, BRN, FDT, FRR and GST), in the western areas (API, IUG, MCD, PRV and YAH), and in a single eastern population (CRC), respectively, and the other populations having a more mixed and less geographically localized genetic composition.

**Figure 2.**
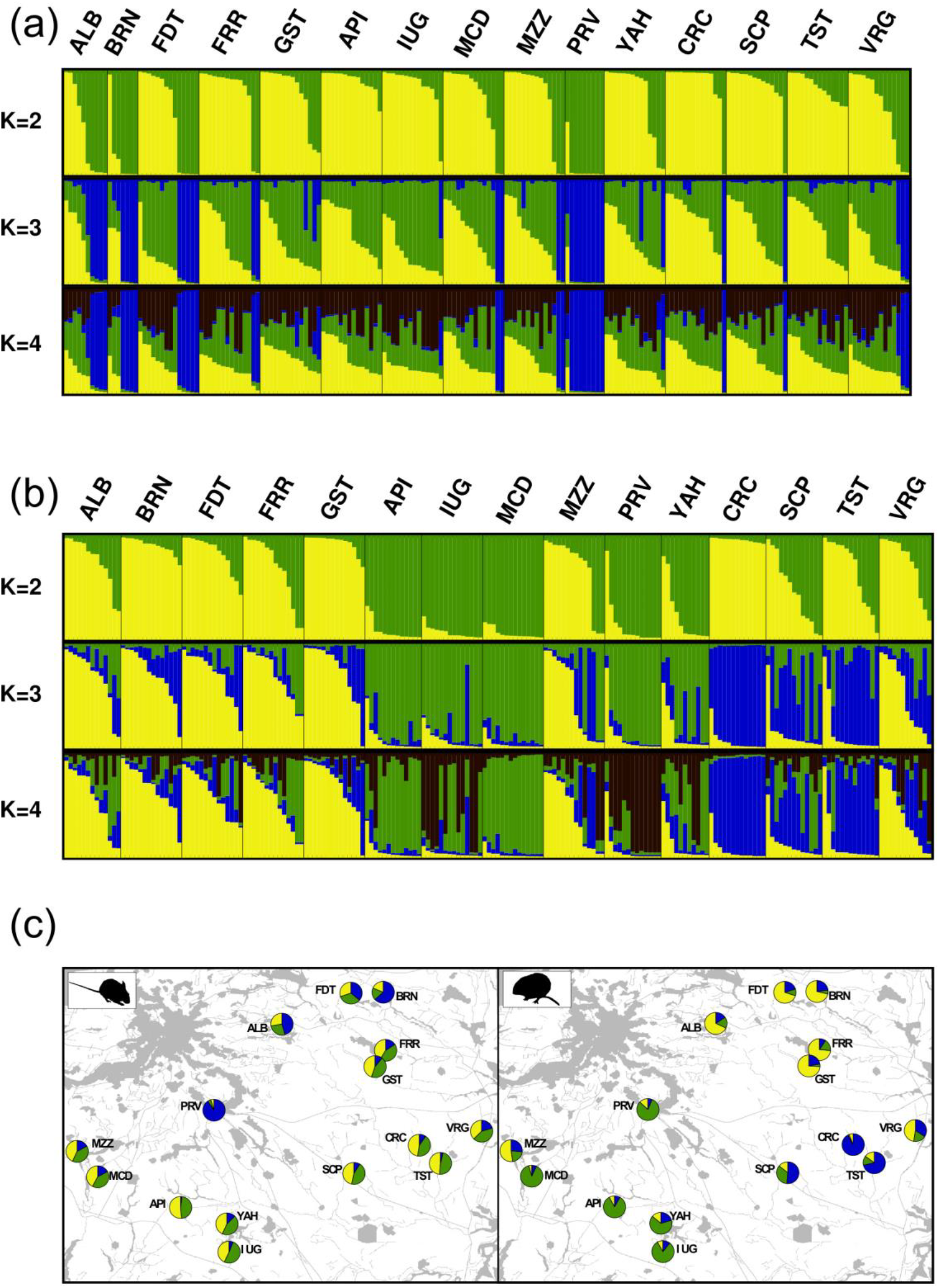
Population assignment test performed with STRUCTURE. Bar plots represent the genetic composition of single individuals (thin vertical columns) from K = 2 to K = 4. A) wood mouse; B) bank vole. (C) Maps of the study area with the genetic composition of each population for K = 3 in the wood mouse (left) and the bank vole (right).

### Visualizing deviation from IBD

The spatial visualization of the geographic areas with higher or lower gene flow compared to IBD expectations is similar in the two species (Fig. 3). The main pattern consists of a central area of reduced gene flow, cantered around PRV, extended only in the bank vole towards the southern and the eastern borders of the region. These branches of reduced migration clearly produce the higher genetic structure observed in the bank vole when compared to the wood mouse, with the latter having a much higher connectivity in most of the areas we considered.

**Figure 3.**
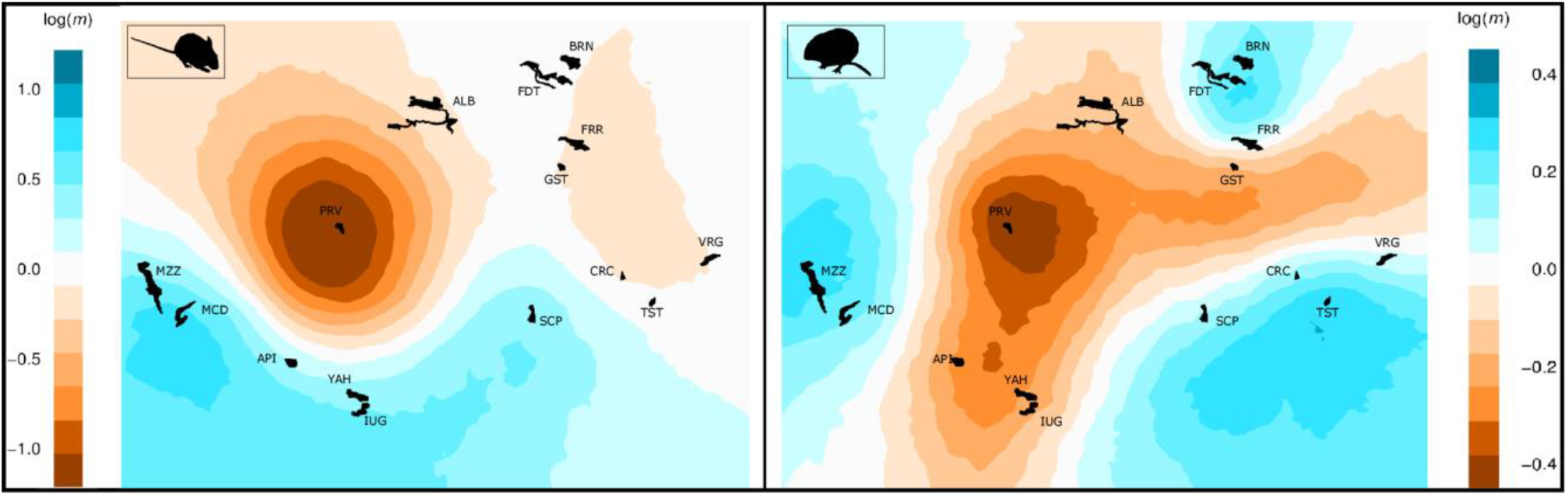
Individual-based EEMS analysis of effective migration rates (m) for the wood mouse (left) and the bank vole (right). The effective migration rate is represented on a log_10_ scale. Areas showing negative values (orange) represent possible barriers to gene-flow while zones with positive values (blue) correspond to places of increased gene-flow, both with respect to the Isolation by Distance background (white). Migration surfaces are averages of 3 runs each with 50, 100, 200, 300, and 400 demes.

### Isolation by resistance

Both the wood mouse and the bank vole populations presented significant patterns of isolation by distance (Supplementary Tables 1-2). However, we also found higher association between pairwise F_ST_ and resistance distance in models including land cover features (Fig 4, Supplementary Tables 1-2). In the wood mouse, the first set of distances (resistance) reached the highest value of *R^2^* when woodland patches presented moderate resistance values (RE =100) with respect to the surrounding environmental feature, while the second set (permeability) highlighted the role of cultivated areas (1/100 of RE_max_) and of the areas comprising and surrounding major roads (1/500 of RE_max_) in facilitating connectivity between different populations. In the bank vole, the resistance scenarios providing the best fit were those implying the highest resistance (RE = 500) for urban areas, whereas woodland and cultivated terrain presented less resistance to gene flow with respect to surrounding land cover (1/500 and 1/100 of RE_max_ respectively). Contrary to the one for the wood mouse (Tab. 3), the cumulative resistance scenario for the bank vole also remained significant once we factored out IBD with partial Mantel tests (*r =* 0.489; p = 0.0384; Mantel test).

**Figure 4.**
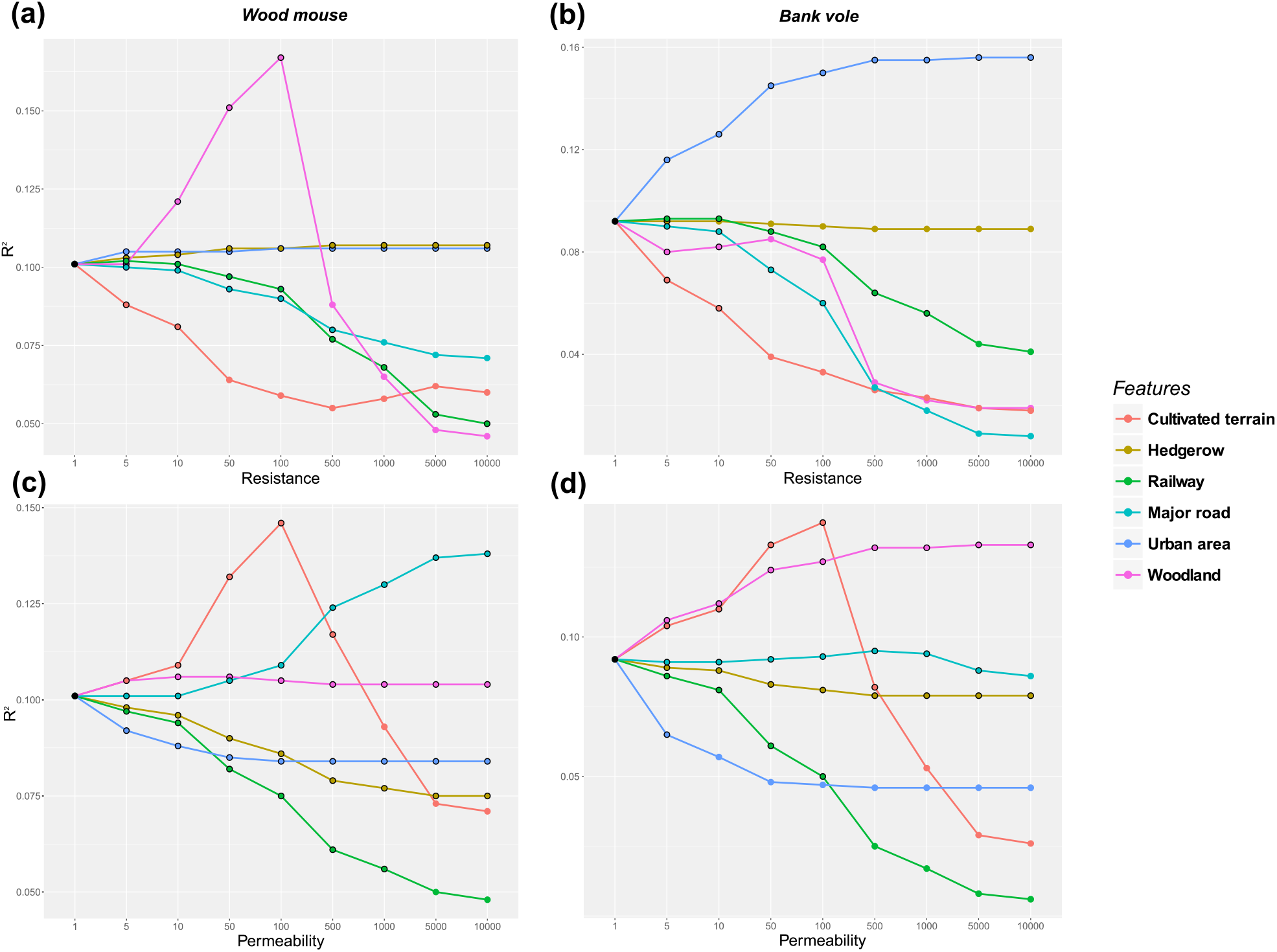
Goodness of fit for models of landscape resistance. Panels show the coefficient of determination (R^2^) for models analysing genetic differentiation (panel A-B: wood mouse; panel C-D: bank vole) in relation to resistance (A, C) and permeability (B, D) distance matrices plotted against resistance values for different landscape features. Circles with black outline showed significant P-values.

**Table 2.** Pairwise F_ST_ distances between sampled populations. Values above diagonal for the bank vole and values below diagonal for the wood mouse. Bold values of F_ST_ indicate significance after Bonferroni correction.

|  | ALB | BRN | FDT | FRR | GST | API | IUG | MCD | MZZ | PRV | YAH | CRC | SCP | TST | VRG |
| --- | --- | --- | --- | --- | --- | --- | --- | --- | --- | --- | --- | --- | --- | --- | --- |
| ALB | - | 0,04 | 0,05 | 0,02 | 0,08 | 0,07 | 0,05 | 0,06 | 0,04 | **0,13** | 0,05 | **0,11** | 0,05 | 0,08 | 0,07 |
| BRN | 0,02 | - | 0,04 | 0,06 | **0,08** | **0,10** | 0,05 | **0,11** | 0,03 | **0,16** | 0,08 | **0,11** | 0,08 | 0,09 | 0,03 |
| FDT | 0,00 | 0,00 | - | 0,01 | 0,06 | **0,12** | **0,07** | **0,09** | 0,04 | **0,11** | **0,08** | **0,13** | 0,07 | **0,09** | 0,05 |
| FRR | 0,00 | 0,00 | 0,01 | - | 0,04 | **0,10** | **0,07** | **0,10** | 0,03 | **0,12** | 0,07 | **0,13** | 0,06 | **0,10** | 0,05 |
| GST | 0,00 | 0,01 | 0,02 | 0,00 | - | **0,13** | **0,08** | **0,12** | 0,05 | **0,15** | 0,07 | **0,10** | 0,07 | 0,07 | 0,04 |
| API | 0,02 | 0,03 | 0,03 | 0,00 | 0,01 | - | 0,05 | **0,09** | **0,09** | **0,17** | 0,03 | **0,11** | 0,03 | **0,09** | **0,11** |
| IUG | 0,00 | 0,02 | 0,02 | -0,01 | 0,02 | 0,01 | - | **0,10** | 0,06 | **0,14** | 0,02 | **0,10** | 0,04 | **0,08** | 0,07 |
| MCD | 0,00 | 0,02 | 0,04 | 0,00 | 0,03 | 0,01 | 0,00 | - | **0,09** | **0,14** | 0,06 | **0,13** | 0,06 | **0,12** | **0,14** |
| MZZ | 0,01 | 0,03 | 0,04 | 0,01 | 0,02 | 0,01 | 0,01 | 0,00 | - | **0,16** | 0,06 | **0,11** | 0,05 | 0,07 | 0,04 |
| PRV | 0,03 | **0,02** | 0,01 | 0,01 | **0,03** | **0,06** | 0,02 | 0,07 | **0,07** | - | **0,15** | **0,21** | 0,13 | **0,22** | **0,17** |
| YAH | 0,01 | 0,00 | 0,00 | -0,01 | 0,00 | 0,00 | -0,02 | 0,02 | 0,00 | **0,05** | - | 0,07 | 0,01 | 0,04 | 0,06 |
| CRC | 0,02 | 0,04 | 0,05 | 0,02 | 0,03 | 0,02 | 0,04 | 0,03 | 0,03 | **0,08** | 0,01 | - | 0,05 | 0,01 | **0,09** |
| SCP | -0,01 | 0,02 | 0,01 | 0,00 | 0,01 | 0,01 | 0,00 | 0,00 | 0,02 | 0,02 | -0,02 | 0,02 | - | 0,04 | 0,05 |
| TST | 0,00 | 0,02 | 0,04 | 0,01 | 0,02 | 0,01 | 0,01 | 0,00 | 0,01 | **0,06** | 0,02 | 0,03 | 0,00 | - | 0,08 |
| VRG | -0,01 | 0,00 | 0,03 | 0,00 | 0,01 | 0,02 | 0,01 | 0,02 | 0,01 | 0,02 | 0,02 | 0,04 | 0,01 | 0,02 | - |

**Table 3.** MMRR and Partial Mantel results for cumulative resistance surfaces. Abbreviation for land cover elements are: cultivated terrain (CT), hedgerow (H), road (Ro), railway (Ra), urban area (Ua) and woodland (W).

|  | Land cover resistance |  |  |  |  |  | MMRR |  |  |  | Partial Mantel |  |
| --- | --- | --- | --- | --- | --- | --- | --- | --- | --- | --- | --- | --- |
| Species | CT | H | Ro | Ra | UA | W | R<sup>2</sup> | β | t | p | r | p |
| Wood mouse | 5 | 10 | 1 | 10 | 10 | 500 | 0.180 | -0.0413 | -3.776 | 0.006 | 0.304 | 0.0773 |
| Bank vole | 5 | 10 | 10 | 10 | 500 | 1 | 0.174 | 0.0219 | 1.645 | 0.001 | 0.489 | 0.0384 |

## Discussion

Our main goal was to investigate the relationship between human-related changes in habitat amount and configuration (i.e., habitat structure), habitat use and genetic structure. We applied the identical sampling scheme within the same fragmented area to two rodent species, the wood mouse and the bank vole. Our major results (see Table 4 for a summary) are that the generalist wood mouse has a population structure much more genetically connected than the forest-specialized bank vole, and cultivated areas facilitate gene flow in both species. Gene flow favoured by cultivated areas likely increases the genetic exchanges in the wood mouse even above the level expected in natural conditions, which appear limited only by woodlands. In the bank vole, cultivated areas possibly act compensating the genetic fragmentation due to the loss of woodland and the increase of urban areas. Overall, we conclude that the difference between these species in their ability to use different habitats is still reflected in the difference between their genetic structure, but this difference is likely to increase if woodlands will be further replaced by urban, but not cultivated areas.

**Table 4.** Concise summary of the major results obtained in the two species.

| Species | Ecology | Overall genetic structure | Main factors limiting gene flow | Main factors favouring gene flow |
| --- | --- | --- | --- | --- |
| Wood mouse | Generalist, found in different habitats | *Expected:* no/low<br>*Observed:* $F_{st}=0.03$; no significant deviation from IBD | Woodland | Cultivated areas; areas around roads |
| Bank vole | Specialist, prefer forests | *Expected:* yes<br>*Observed:* $F_{st}=0.08$; significant deviation from IBD | Urban areas | Cultivated areas; woodland |

### Genetic diversity

Habitat fragmentation did not produce a detectable loss of genetic variation in two species. Levels of diversity in different populations are comparable to those reported for other rodent species^40, 71–73^. When the global genetic divergence between populations is analyzed, the wood mouse shows much weaker population structure than the bank vole. This pattern is expected considering that, at a short geographic scale (distances <30 km), genetic structure is commonly found only in rodents with a specialized ecological niche^73–79^.

With the exclusion of the population sampled in PRV (see below), the wood mouse appears rather homogenous at this geographic scale, indicating that gene flow is not prevented by the human-induced fragmentation of their natural habitat. This result reflects the enormous capacity of adaptation and mobility in this species, which can be found in all types of forests and even in cultivated fields in some periods of the year^80–82^. On the other hand, populations of the bank vole sampled in the same patches showed the presence of a significant genetic differentiation with a lower degree of genetic admixture and higher F_ST_ values. Similar studies on bank vole confirmed that there is a significant reduction of gene flow already at geographical distance of about 8 km^83^, and that environmental features, such as seasonal temperature variations, can contribute in a decisive way in increasing the genetic structure of this species^84^.

### Spatial patterns of gene-flow

Isolation by distance was significant, indicating that geographic distance is an important factor for both species. An additional shared feature appears the isolation of PRV in all the analyses, supporting the hypothesis that individuals in both species have some difficulty to reach this area. This result may be related to the fact that woodland and urban areas are highly diffused around PRV, and the IBR analysis suggested that woodland acts as a barrier for the wood mouse whereas urban areas act as a barrier for the bank vole.

The relevance of woodland as a barrier for the wood mouse can be explained by the competition with the forest specialist bank vole or/and with the congeneric species *Apodemus flavicollis*, as shown by empirical studies of the strength of interspecific competition in shaping small mammal communities in fragmented landscapes^32^.

Additional areas of enhanced or reduced gene flow, in comparison with the isolation by distance pattern in the background, were found for the bank vole. Specifically, three main areas showed gene flow higher than expected, corresponding to western, eastern and northern patches. Barriers separating them are composed of a mix of different environmental features, but the IBR modelling suggests that urban areas play the major role.

Finally, a few general comments on the results provided by the IBR analyses are needed. Railways and roads (never wider than 10 meters in this area) cannot be considered as barriers to the dispersal of these species, consistently with previous studies^71,76^. Indeed, roads appear as a factor that favours gene flow in the wood mouse. This may be because, for this species, the size of the roads present in the study area should not be considered as a barrier and/or that roads, in the environmental matrix, were included in (or surrounded by) a suitable ground. Similarly, cultivated fields do not limit dispersal, but may even play a role as corridors^85^. The only anthropogenic factor that seems to negatively affect the dispersal pattern (only in the bank vole) is the presence of urban areas. Clearly, if woodlands will be further reduced by urbanization, genetic fragmentation could become an issue for the bank vole, but not for the wood mouse.

### Conclusions and implications for conservation

Overall, the results of this research show that, despite extensive habitat changes due to human activities, levels of genetic variation are quite high in both species, and their difference in the dispersal abilities is still reflected in the difference of genetic structure. The wood mouse, a generalist species with high dispersal ability, shows in fact higher genetic connectivity than the bank vole, which is a less mobile species closely linked to woodland areas. Nevertheless, we found also that cultivated fields and urban areas modifies the natural dispersion patterns in both species, probably in a way that will, in the future, increase the difference between their genetic structure. Our study supports the view that patterns of gene flow can be differently affected, even in related and sympatric species, by the same changes of land use. Locally, this implies that future monitoring efforts should prioritize the bank vole, the species with the highest genetic structure where genetic fragmentation is more likely to increase due to urbanization. More in general, we argue that predicting the genetic impact of habitat fragmentation using single model species may be misleading.

## Acknowledgments

RB’s stay at the University of Potsdam was supported by the Erasmus Placement grant (UU-ER/2010/010744, M.S.) from the European Commission Life Long Learning programme. RB thanks Ralph Tiedemann for his hospitality at the Unit of Evolutionary Biology/Systematic Zoology (Institute of Biochemistry and Biology, University of Potsdam) during the laboratory analysis. We thank all students that helped us with fieldwork.

## Author Contributions

R.B., A.M., V.K. and G.B. conceived and designed the study. A.M. and G.S. obtained the samples. R.B. and K.H. carried out the laboratory work. R.B. and A.B. analysed the data. R.B., A.B. and G.B. wrote the manuscript, with contributions from all the authors.

